# How the West(ern) Was Won: Solutions for Immunoblotting Large and Small Proteins

**DOI:** 10.1101/2022.03.23.485494

**Authors:** Paula Llabata, Pere Llinàs-Arias

## Abstract

Relative protein quantification is a well-established technique in the vast majority of the molecular biology laboratories. However, western blot standard protocols may be unable to detect certain proteins depending on their size. When the protein of interest is out of 10-250 KDa range, its migration through the gel or transfer to the membrane are compromised, thus making its detection difficult. Here we present a set of modifications of the standard working procedure for western blotting based on the experience working with Small VCP Interacting Protein (SVIP) and Max-Gene Associated Protein (MGA), whose molecular weight are 8 and 350 kDa, respectively. We expect that these adaptations may help researchers to improve their experiments in a cost-effective manner.

## 1. INTRODUCTION

During this century, the rise of high throughput techniques has allowed an unprecedented advance in our knowledge about molecular biology [1, 2]. The implementation of “omics” (including genomics, proteomics, transcriptomics and metabolomics, among others) and its combination has contributed to a better understanding of different diseases describing new biological networks, connecting the dots and finding out its clinical relevance [3]. Although the relevance of these techniques, the results derived from these studies must be validated through different classic methods. In addition, many studies are focused on a single pathway, where a simpler and cost-effective method is needed. Regarding to the study of proteins, the standard method is western blot. The Introduction section should include the background and aims of the research in a comprehensive manner.

Western blot is a well stablished technique which is performed in the vast majority of molecular biology laboratories to determine the relative protein abundance among different experimental conditions. After antibody optimization, a standard protocol usually works for a large proportion of proteins, which are comprised between 10-250 kDa (Fig. 1A). However, these procedures are not valid when the target protein size is not included in the abovementioned range.

**Fig. (1).**
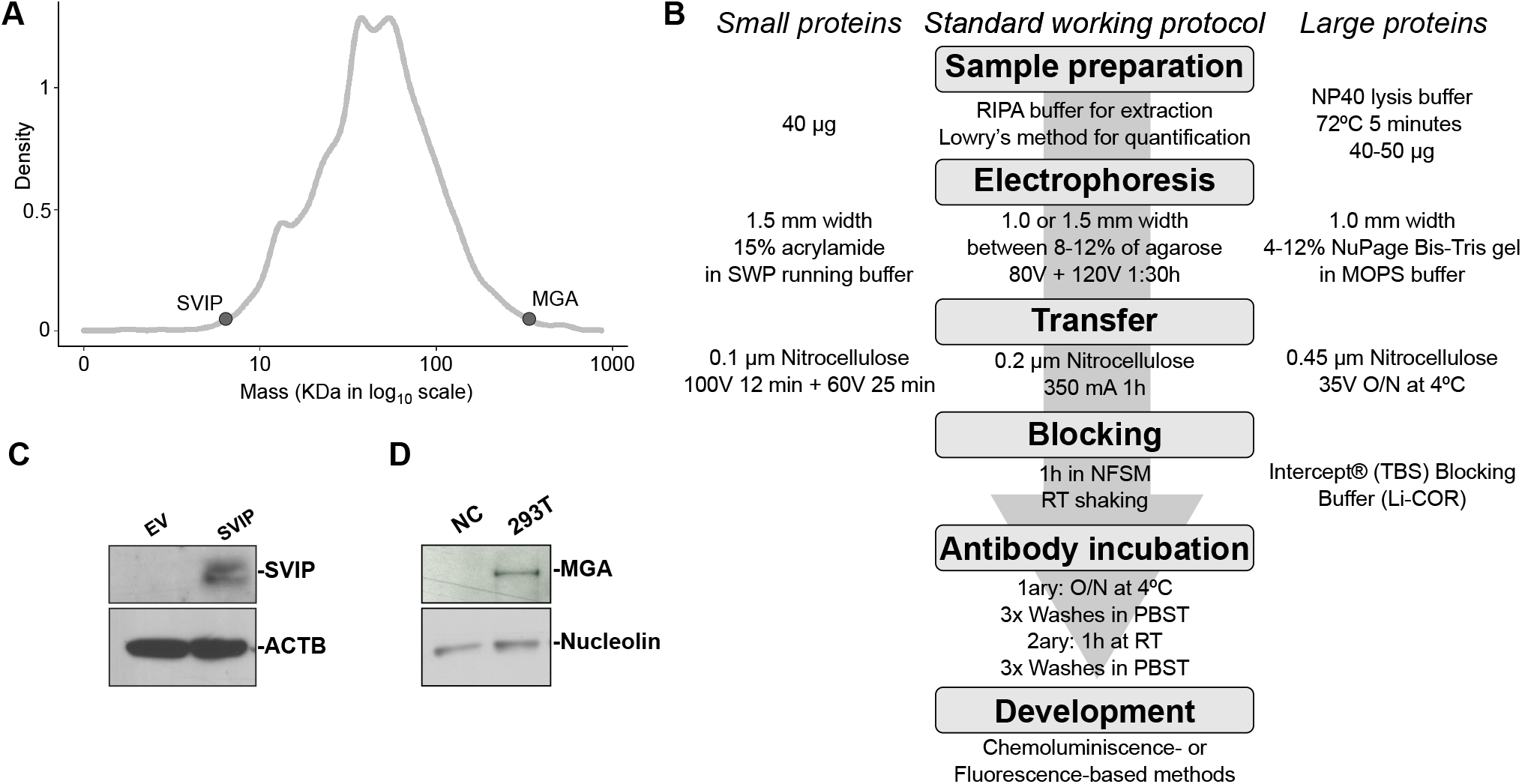
**(A)** Density plot of human proteome based on Uniprot data. Protein mass in logarithmic scale is represented, and SVIP and MGA appear highlighted. (**B**) Summary workflow for western blot on standard conditions and their adaptations for small and large proteins. Examples of western blots against (**C**) SVIP and (**D**) MGA performed modifying the standard protocol as shown in B. SVIP western blot was performed in BB30-HNC cell line either transfected with pLVX-ZSgreen empty vector (EV) or containing SVIP CDS. MGA western blot was performed in LouNH91 as negative control (NC) and HEK 293T transfected with pLVX-ZSgreen containing MGA CDS. NFSM: Non-fat skim milk.

Here we describe a set of modifications from the Standard Work Procedure (SWP) of our laboratories to perform Western blots to detect either small or large proteins. These changes were used in different studies involving Small VCP Interacting Protein (SVIP) [4] and Max-Gene Associated Protein (MGA) [5, 6], and they led to overcome the technical issues related with the above-mentioned targets (Fig. 1B).

## 2. MATERIALS AND METHOD

### 2.1. Sample preparation

Adherent cells are collected using a scrapper after washing cells in PBS twice followed by centrifugation (2000 rpm, 2 min, 4°C). Suspension cells are directly collected through centrifugation (2000 rpm, 2 min, 4°C), washed and centrifuged again. Frozen tissue samples – derived from orthotopic tumours, for instance – are mashed in a pestle in presence of liquid nitrogen to prevent its degradation. Once the sample is homogenised, protein extract is obtained following the next step.

Total protein extracts are obtained using RIPA buffer (20 mM Tris-HCl [pH 7.5], 150 mM NaCl, 1 mM Na2EDTA, 1 mM EGTA, 1% NP-40) buffer (100 μL for 2·106 cells, approximately). Then, cell lysates are sonicated during 5 min and gently mixed in the rotator wheel during 15 minutes at 4°C. After that, tubes are centrifuged 15 min at 13000 rpm.

Collected supernatants are quantified through the Lowry method [7], using the DC™ Protein Assay Kit II (BioRad, 5000112). Then, samples are prepared for electrophoresis: quantified protein extracts are diluted to equal concentration (around 10-30 μg depending on the availability), mixed with Laemmli buffer (final concentration: 62.5 mM Tris-HCl pH 6.8, 25% glycerol, 2% SDS, 0.01% Bromophenol Blue, 5% β-mercaptoethanol) and boiled during 5 minutes at 95°C.

### 2.1. Sample preparation

Tris glycine-SDS-Polyacrylamide gels are prepared in empty cassettes (Thermofisher, NC2015) pouring the resolving and stacking mixes. Resolving mix must be firstly prepared and rapidly dumped into the empty cassettes. This mix determines the gel density, and allows a proper separation depending on your protein of interest. Different recipes varying % acrylamide are included in **Table 1**. After pouring the resolving mix, 2ml of isopropanol is added on the top of the gel. This step will accelerate acrylamide polymerization process. Once the resolving mix has polymerized and isopropanol has been removed, the stacking mix can be prepared and loaded into the cassettes, including a comb with the desired number of wells.

**Table 1:**
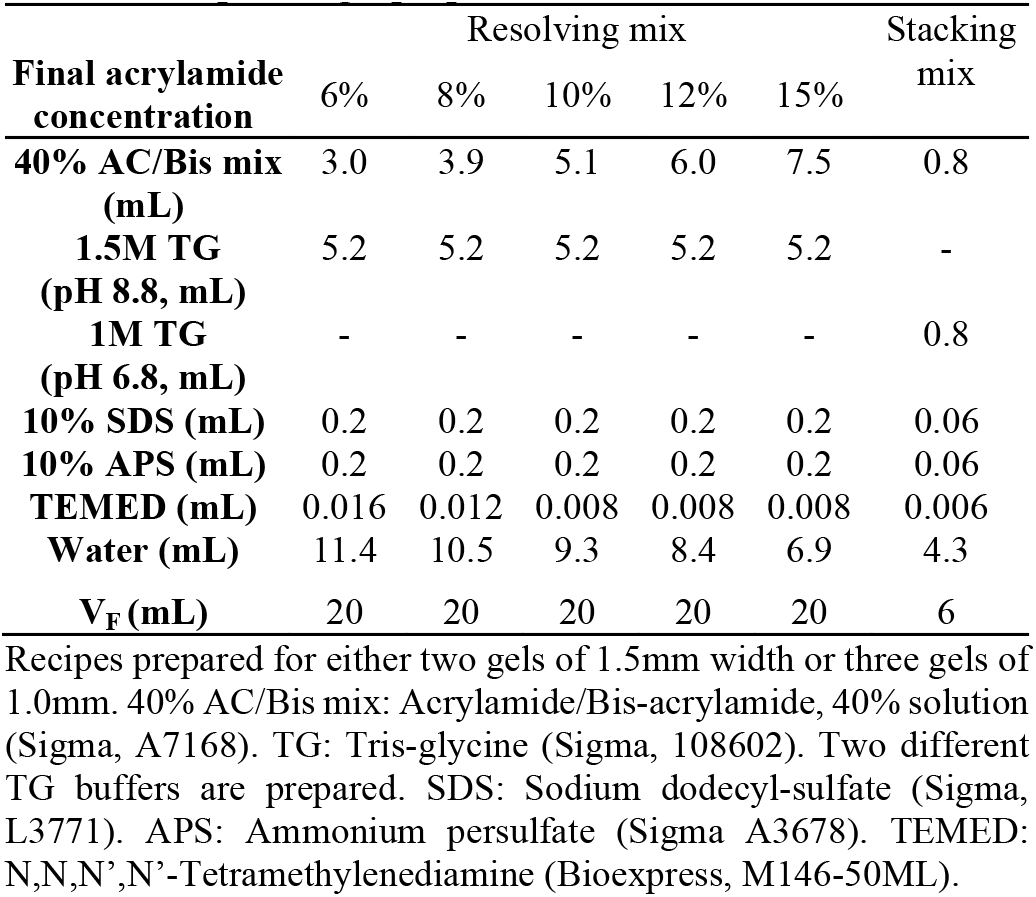
Recipes for gel preparation.

The polymerized gel is set into a mini gel tank (A25977, Thermofisher) and filled with running buffer (25 mM Tris, 200 mM glycine, pH 8.5). After removing the comb, samples and the protein marker are loaded in the gel where the electrophoresis takes place. This step may force the proteins to migrate depending on their size. Electrophoresis is performed in two steps maintaining constant voltage: while protein extracts travel through the stacking phase, 80 V are recommended. Once the sample has entered into the resolving phase, voltage can be increased up to 120 V. Typically, electrophoresis is stopped when the dye front reaches the bottom of the gel, which spends 90 to 120 minutes.

After electrophoresis, the separated protein mixtures are transferred to 0.2 μm nitrocellulose membranes (GE10600001, Amersham) using Wet/Tank blotting systems (1703930, Biorad). Cassettes containing the gel and the membrane, as well as filter papers and sponges undergo to a constant amperage of 350 mA during 1h in transfer buffer.

Under these conditions, proteins migrate from gel to membrane. Transfer buffer is based on running buffer supplemented with 20% methanol. This step generates heat that may compromise the experiment. For that reason, the Wet/Tank must be cooled during the whole process using cooling unit inside the tank, covering it with ice, or performing the transfer step in a cold room.

### 2.3. Blocking, antibody incubation and development

The membrane, which contains the separated proteins, is blocked using 5% Non-Fat Skim Milk (NFSM; 232100, Becton Dickinson) in phosphate buffered saline (PBS) containing 0.01% Tween 20 (PBST; 85113, Thermofisher) during at least 1h at room temperature on a shaker. Then, the membranes covered by a dilution of the primary antibody in either 5% NFSM or 5% BSA (antibody dilution is indicated by the manufacturer). This incubation is performed over night at 4°C on a shaker.

The following day, the primary antibody mixture is removed from the membrane. The membrane is washed at least three times using PBST. Washes take place at room temperature during five minutes each. Then, the membrane is incubated with a secondary antibody during 1h at room temperature. Secondary antibody conjugation depends on the developing method. IRDye antibodies (Li-COR) are used for an Odyssey Imaging System (Li-COR), whereas HRP-conjugated secondary antibodies are based on chemoluminiscence (WBLUC0100, WBLUF0100, WBLUR0100, all from Merck Millipore), which is detected by light-sensitive films (Hyperfilm ECL, 28-9068-37, GE HEALTHCARE) and further developed using a chemical system (Ilford).

## 3. SIGNIFICANT CHANGES FOR SMALL PROTEIN DETECTION

SVIP is an 8 KDa protein composed by 78 amino acids [4]. The detection of this protein is challenging, since it is not detected following the SWP even after its ectopic overexpression in a cancer cell line (data not shown). Samples containing 40μg are prepared following the SWP. For optimal separation, protein extracts are loaded into a 15% acrylamide SDS-PAGE gel with 1.5 mm of width. These gels are prepared on empty cassettes (Thermo Scientific, NC2015).

Electrophoresis starts at 80V. Once the front has entered into the resolving part of the gel, the voltage is increased to 140V during 75 min. Then, proteins are transferred into a 0.1 μm pore size nitrocellulose membrane. Wet transference is performed at constant voltage, 100V during 12 minutes followed by 60V during 25 minutes. These steps allow SVIP detection as shown in Fig. 1C.

## 4. SIGNIFICANT CHANGES FOR LARGE PROTEIN DETECTION

MGA is a 350 KDa estimated weight protein composed by 3,065 amino acids [5, 6]. The detection of this protein is arduous, since it cannot be detected following the SWP even after its ectopic overexpression in a cancer cell line.

For improved immunoblotting, protein samples are lysed in NP-40 lysis buffer (1% NP-40, 150 mM NaCl, 50 mM Tris-HCl pH 8.0) supplemented with protease and phosphatase inhibitor (Thermo Scientific, 1861281), followed by denaturation in LDS sample buffer (Thermo Fisher) supplemented with 20 mM DTT, since the traditional homemade loading buffers are not suitable due to its high amount of SDS. Protein extracts are boiled at 72°C during 5 minutes. Protein standard should also be considered, since most of the protein standards do not include molecular weights greater than 250 KDa. For this purpose, a protein marker reagent containing molecular weights up to 460 KDa (HiMark™ Pre-stained Protein Standard, ThermoFisher) is selected.

Electrophoresis is performed in 4-12% NuPage Bis-Tris 1.0 mm width gel in MOPS SDS running buffer (Novex™, Life technologies). Protein transfer is the biggest challenge for high molecular weighted proteins. For this case, the amount of input protein for wester-blot should also been adjusted. Any amount below 30μg may lead to undetectable band after transfer. A range of 40-50μg per well is loaded. Proteins are then transferred to a nitrocellulose membrane (0.45 μm, Amersham™ Protran™ NC) in Tris-Glycine Transfer Buffer (Thermo Scientific) supplemented with 10% methanol overnight at 4°C at constant voltage of 35V. As mentioned above, the amount of protein needed for detectable band after transference is higher than the required in the SWP. Moreover, the exposure time for obtaining a clear band may be also higher, therefore, development of western-blot images is also challenging. For these reasons, chemoluminiscence method was discarded in order to avoid membrane burning. Instead, images are taken with the Odyssey® Imaging Systems instrument whose developing method is based on fluorescence.

## CONCLUSION

The abovementioned changes led to overcome technical issues and efficiently develop protein analysis. We expect that these steps may help other researchers to detect small and large proteins, which in turn may contribute to increase knowledge of molecular biology.

## AUTHORS’ CONTRIBUTIONS

Paula Llabata and Pere Llinàs-Arias performed the optimization of MGA and SVIP western blots, respectively. Both conceived and wrote the manuscript.

## ACKNOWLEDGEMENTS

Western blot experiments of SVIP and MGA were performed upon the direction of Manel Esteller and Montse Sánchez-Céspedes, respectively. Paula López-Serra and Lida Rosselló-Tortella also participated in SVIP western blot optimization.

## Notes

### Competing Interest Statement

The authors have declared no competing interest.

